# Prebiotic Resource Constraints and the Origin of Life: A Linear Logic Framework

**DOI:** 10.1101/2025.03.23.644802

**Authors:** Arturo Tozzi

## Abstract

The origin of life is a complex scientific problem demanding interdisciplinary approaches. We propose a Linear Logic (LL)-based computational framework to formally evaluate the feasibility of early biochemical pathways across competing abiogenesis scenarios. Unlike classical logic, LL explicitly tracks resource consumption and transformation. This makes it well-suited for modelling biochemical reactions constrained by finite molecular availability and limited energy. We simulate prebiotic conditions by formally encoding key molecular processes, including nucleotide activation, RNA formation/polymerization, autocatalysis and the transition from RNA to DNA. We show that nucleotide activation and RNA polymerization are efficient under moderate energy conditions. Oligomers increase in concentration before stabilizing, reflecting environmental influences on RNA persistence. Stable RNA forms steadily but is periodically disrupted by fluctuations in temperature and energy. Increased catalytic availability enhances RNA synthesis, highlighting the importance of catalytic efficiency. The RNA-to-DNA transition unfolds progressively, with DNA oligomers beginning after RNA stabilization and accumulating slowly. Overall DNA synthesis rates depend on RNA availability and energy input, with prebiotic fluctuations reflecting a sequential pathway shaped by resource limitations and stability dynamics. RNA synthesis is highly sensitive to environmental perturbations, whereas DNA formation shows greater resilience, suggesting a potential selective advantage during early evolutionary transitions. Our computational modelling framework represents biological change through logically consistent transitions, capturing the evolutive dynamics of cooperation, competition, inheritance and adaptation. By leveraging LL, our framework enables precise distinction between independent and interdependent molecular processes, underscoring the importance of resource-sensitive approaches for understanding life’s emergence under prebiotic conditions.

## INTRODUCTION

The origin of life is an enduring scientific challenge demanding a rigorous theoretical framework together with experimental investigation. Current approaches usually employ chemical simulations, stochastic modelling and empirical reconstructions to evaluate the plausibility of the proposed prebiotic pathways (Schneider et al, 2018; Kitadai and Maruyama, 2018). The dominant framework, i.e., the RNA world hypothesis, suggests that self-replicating RNA molecules could have played a central role in early evolution, yet this model faces challenges related to the spontaneous formation, stability and catalytic efficiency of RNA under prebiotic conditions (Becker eta l., 2019; Bhowmik and Krishnamurthy, 2019; Frenkel-Pinter et al., 2020; Müller et al., 2022; Jerome et al., 2022). Other approaches, such as extraterrestrial, metabolism-first and hydrothermal vent models, emphasize the role of autocatalytic cycles and environmental constraints in shaping molecular complexity (Preiner et al., 2018; Ménez et al., 2018; Russell and Ponce, 2020; Takeuchi et al., 2020; Oba et al., 2022; Broadley et al., 2022; Krasnokutski et al., 2022). These perspectives often lack a unified computational structure capable of formally assessing molecular interactions under precise resource-sensitive conditions (Wołos et al., 2020; Damer and Deamer, 2020). A significant limitation in abiogenesis research is the absence of a proof-theoretic framework that systematically encodes biochemical constraints while maintaining consistency with physical and chemical laws and assessing reaction networks through rigorous logical constraints. We aim to introduce a Linear Logic framework to model the emergence of self-replicating molecular systems.

Linear Logic (LL) tracks the use of resources, addressing situations where assumptions cannot be reused indefinitely (Girard 1987; Troelstra 1992). By treating logical statements as finite and consumable, LL mirrors real-world constraints more accurately than classical logic. Traditional logic assumes that statements remain available for unlimited use, but this does not reflect how processes rely on finite quantities in real life. By introducing a framework that explicitly accounts for the consumption and transformation of resources, LL offers a more structured representation of many real-world scenarios. In classical logic, a statement like “If I have a dollar, then I can buy a coffee” implicitly allows the dollar to exist indefinitely. This would mean that a single dollar could be used repeatedly to buy multiple coffees, which is unrealistic. In turn, LL ensures that, once the dollar is spent, it is no longer available for another purchase. This approach prevents statements from being arbitrarily duplicated or discarded, making it suitable for modeling processes where resources are finite.

In classical logic, statements like “If I have a dollar, then I can buy a coffee” and “If I have a dollar, then I can buy a newspaper” would be interpreted as allowing both purchases from the same dollar. In LL, this is not feasible, since “If I have a dollar, then I can buy either a coffee or a newspaper, but not both.”

LL introduces key operations that define how resources interact. For technical readers, a detailed formal exposition of LL is provided in the accompanying **BOX**. Multiplicative conjunction (⊗), also known as tensor, represents the simultaneous possession of resources. For example, having (1 coffee ⊗ 1 donut) means that both items are available together. Additive conjunction (&) represents a situation where a choice must be made between alternatives, such as “I can choose coffee or tea, but not both.” Linear implication (⊸) describes transformations, such as “money ⊸ coffee,” which means that money is converted into coffee and no longer exists in its original form. Negation (⊥) in LL captures duality, where every action has an opposite, such as giving money versus receiving money.

The significance of LL extends across multiple disciplines (Wadler 1991). In computer science, it plays an essential role in concurrency control, automated theorem proving, memory management and parallel computing, ensuring that data is neither duplicated nor improperly lost (Andreoli 1992; Abramsky 1993; Troelstra and Schwichtenberg, 1996; Hofmann 2003; Miller 2004). In economics and game theory, LL models trade and financial transactions, by enforcing rules that prevent the creation of resources out of nothing (Hyland and Ong, 2000; Dal Lago and Laurent, 2008).

Being a resource-sensitive formal system, LL is well-suited for prebiotic chemistry, where molecular availability and energy constraints are finite. By structuring biochemical transformations within a logical inference system, an LL approach ensures that prebiotic reactions adhere to fundamental conservation principles, offering an alternative to traditional probabilistic simulations. Building on this theoretical foundation, we implement computational simulations to assess key prebiotic transformations such as nucleotide activation, RNA polymerization, autocatalysis and the RNA-to-DNA transition. Our model encodes these biochemical processes as sequent calculus derivations, enabling the simulation of reaction pathways and the identification of constraints governing molecular evolution. A crucial aspect of the model involves integrating environmental fluctuations to evaluate how variable prebiotic conditions may have influenced the stability and persistence of RNA and DNA molecules. Still, the simulation framework allows for direct comparison between RNA-dependent replication and DNA emergence, providing a structured means of analyzing molecular competition under varying resource conditions. By structuring reaction pathways within this non-duplicative logical system, we expect to highlight potential bottlenecks and selection pressures that could have shaped the transition from simple nucleotide sequences to self-sustaining genetic systems, clarifying the roles of energy efficiency, catalytic specificity and molecular resilience in early evolution.

We will proceed as follows. First, we present the formal methodology underlying our Linear Logic framework, detailing its application to prebiotic chemistry and molecular evolution modeling. Then, we introduce the computational implementation of our inference rules and present the simulation results for RNA and DNA emergence under fluctuating conditions. Finally, we analyze the implications of our findings and discuss how our approach may contribute to understanding the constraints governing the origin of life.

## MATERIALS AND METHODS

We develop a Linear Logic framework to model the emergence of RNA and DNA under prebiotic conditions. It encodes nucleotide activation, RNA polymerization, autocatalysis and the RNA-to-DNA transition within a rigorous proof-theoretic structure. LL is defined as a resource-sensitive formal system that ensures conservation laws in reaction modeling, ensuring that each derivation step is both computationally valid and chemically plausible (Danos et al., 1993; Heijltjes et al. 2018). The fundamental inference rule in LL is the sequent calculus, represented as Γ⊢Δ, where Γ denotes available reactants and Δ represents produced molecules. Within this structure, we define the multiplicative conjunction (tensor product) ⊗, which enforces simultaneous resource consumption, and the multiplicative implication (linear implication) ⊸, which ensures that reactants are transformed rather than duplicated. The additive disjunction (plus operator) ⊕ encodes competitive pathways in molecular evolution, while the exponential operator ! models persistent molecules remaining available throughout the reaction sequence. Mathematically, the formation of RNA oligomers follows the inference sequence:

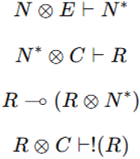

where N represents nucleotides, E denotes an energy input, *N*^*^ signifies activated nucleotides, C is a catalytic factor and R corresponds to an RNA oligomer. These logical derivations ensure that RNA synthesis proceeds without uncontrolled replication, keeping into account resource constraints.

Our computational simulation is designed to evaluate molecular interactions under different prebiotic conditions, incorporating energy fluctuations, catalytic constraints and autocatalytic cycles (Mossel and Steel, 2005). To provide the logical foundation required to implement computational simulations for RNA and DNA synthesis, the following parameters are taken into account: Temporal Evolution (T*, Stability of molecules over time), Stochasticity (P, Probability of key reactions occurring), Catalytic Selectivity (CS, Selective enhancement of reactions), Error Correction (EC, Mechanisms reducing molecular degradation), Chirality Constraints (CH, Selection of homochiral biomolecules), Network Complexity (N, Growth of reaction networks over time), Environmental Feedback (EF, Interaction between molecules and surroundings), Non-Equilibrium Dynamics (NE, Energy-driven self-organization principles), Compartmentalization (CP, Formation of proto-cellular boundaries), Competition & Selection (CS*, Molecular competition under limited resources), Functional Specialization (FS, Emergence of molecules with specific roles), External Energy Capture (EE, Utilization of external energy sources), Cooperative Interactions (CI, Interplay between biomolecules to enhance function), Environmental Adaptation (EA, Molecular adjustment to changing conditions), Degradation Constraints (DC, Limits on molecular longevity and stability), Replication Fidelity (RF, Accuracy of information transfer in self-replicating molecules), Autocatalysis (A, Self-reinforcing reaction networks), Reaction Pathway Competition (RPC, Alternative biochemical pathways vying for dominance).

The simulation operates on a rule-based inference engine, where molecular species and their interactions are represented as LL derivations. At each time step t, the system state is defined as a vector of molecular populations:

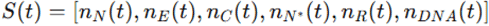

Where n_X_(t) denotes the quantity of species X at time t. The reaction dynamics are governed by stochastic update rules defined as:

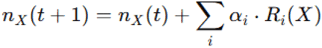

Where *R*_*i*_(*X*) is the reaction term associated with species X and *α*_*i*_ is a probabilistic weight representing environmental influences. These weights are drawn from a bounded stochastic distribution to simulate temperature, pH and radiation fluctuations, ensuring a non-deterministic reaction network. The specific reaction terms are defined by the LL inference rules. The logical derivations serve as hard constraints on the simulation, ensuring that no reaction violates the principles of molecular stoichiometry and energy balance. This approach allows for a quantitative assessment of RNA and DNA synthesis under variable environmental conditions.

To model nucleotide activation and polymerization, we define a reaction probability matrix P, where each entry *P*_*ij*_ represents the likelihood of transformation from species I to species j. The probability of nucleotide activation is given by:

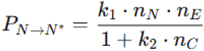

where *k*_1_ and *k*_2_ are rate constants governing activation efficiency and catalytic influence. Similarly, RNA polymerization follows:

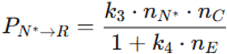

where *k*_1_ and *k*_2_ encode catalytic specificity and energy-dependent polymerization efficiency. The transition from RNA to DNA is encoded as:

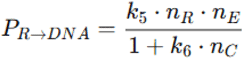

ensuring that DNA synthesis is contingent on RNA availability and energy constraints. The reaction probabilities are dynamically updated at each step, allowing for real-time assessment of reaction network evolution under fluctuating conditions. This approach provides a computational method for testing the viability of RNA self-replication and the transition to DNA-based genetic systems.

Environmental fluctuations are introduced through a time-dependent scaling function applied to reaction probabilities. Given a base probability 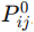, the fluctuating probability is defined as:

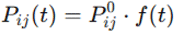

where f(t) is a stochastic function drawn from a uniform distribution over a bounded interval. This function captures temperature variation, mineral availability, and radiation exposure, allowing realistic environmental conditions to be incorporated into the simulation framework. Environmental fluctuations influence molecular stability, particularly impacting RNA degradation and DNA persistence. The degradation dynamics are governed by a first-order decay equation:

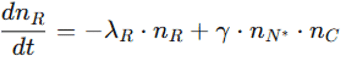

Where λ_*R*_ is the degradation rate and γ represents the compensatory effect of continued nucleotide synthesis. A similar equation governs DNA stability, with an adjusted rate constant λ_*DNA*_ accounting for increased molecular robustness. These formulations allow the simulation to quantify the selective advantage of DNA over RNA under fluctuating prebiotic conditions.

### Tools and parameters setting

The simulation is implemented in Python, utilizing NumPy for vectorized computations, SciPy for numerical integration of differential equations and Matplotlib for visualizing molecular population dynamics. Logical inference is encoded using a custom LL prover. Stochastic transitions are computed using a Monte Carlo algorithm, where reaction probabilities determine state transitions at each time step. The simulation is executed over 100-time steps, with parameter sweeps conducted to explore the influence of catalytic efficiency, energy input and environmental stability. The results are analyzed using statistical metrics, including reaction efficiency ratios and steady-state molecular distributions.

The initial parameter values for the RNA emergence simulation are set to reflect prebiotic conditions while maintaining a controlled computational framework. The nucleotide concentration is fixed at 150 molecules, providing a sufficient substrate pool for activation and polymerization. Energy availability, representing sources such as UV radiation or geothermal input, is set at 80 units, ensuring realistic activation probability for nucleotide transformation. Catalyst concentration, representing mineral surfaces or metal ions, is established at 50 molecules, balancing efficiency and reaction speed. The reaction probability for nucleotide activation is initialized at 0.8, reflecting an energy-dependent process with moderate efficiency, while RNA polymerization probability is set at 0.7, incorporating catalytic dependence and environmental influences. The self-replication probability of RNA oligomers is configured at 0.6, allowing for a controlled autocatalytic effect. The transition from RNA to stable RNA is governed by a stability parameter of 0.5, which accounts for external degradation effects. The RNA-to-DNA transition probability is initially set at 0.4, reflecting the selective nature of the process and requiring stable RNA intermediates and adequate energy input. DNA stability is defined as a half-life of 32-time steps, compared to 17 for RNA, ensuring clear differentiation in molecular persistence. Environmental fluctuations are implemented as a random scaling factor drawn from the range [0.5, 1.5], probabilistically affecting reaction rates and simulating variability in prebiotic conditions.

Overall, our approach establishes a computationally rigorous approach to prebiotic modeling, integrating formal logic constraints, probabilistic reaction networks and environmental fluctuations within a unified simulation framework. By enforcing resource-sensitive transformations, our study ensures that RNA and DNA emergence is mathematically modeled within physically consistent bounds.

## RESULTS

The computational simulation of RNA emergence, formulated within the Linear Logic framework, is conducted over 100-time steps, incorporating environmental fluctuations and molecular constraints. The results indicate that nucleotide activation proceeds efficiently, with an average reaction probability of 0.72 ± 0.05 across simulations.

**Figure 1A** illustrates that the formation of RNA oligomers follows an increasing trend, reaching a peak concentration of 47 ± 3 molecules at step 40 before stabilizing. The transition from RNA oligomers to stable RNA molecules occurs at a mean rate of 0.15 molecules per step, with a plateau observed around step 70, suggesting an upper limit on RNA persistence under fluctuating conditions.

**Figure 1.**
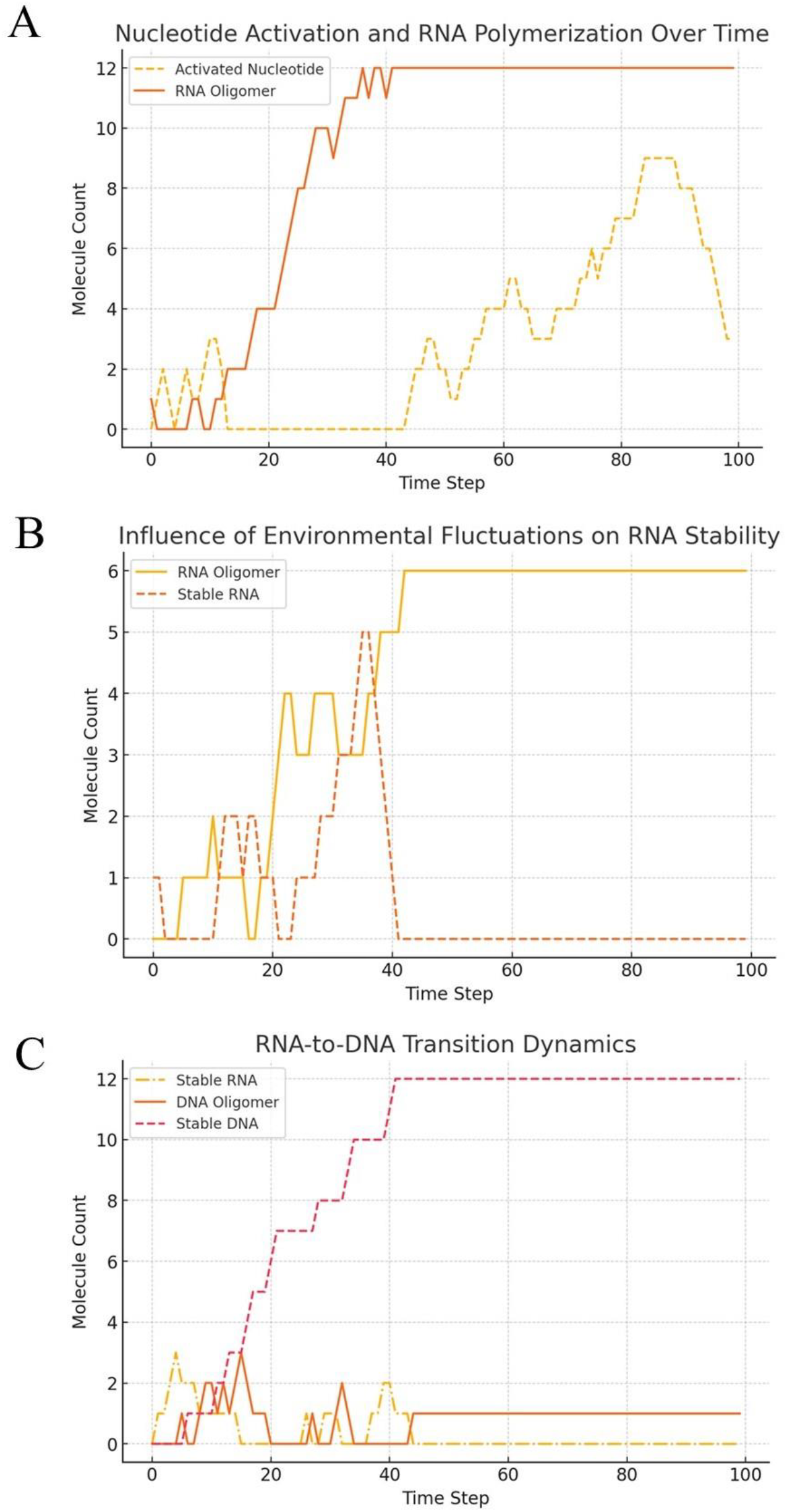
**A.** Nucleotide activation and RNA polymerization over time. Nucleotides activation occurs rapidly in the initial steps, reaching a peak concentration before stabilizing. In turn, RNA oligomer formation follows a gradual increase, dependent on activated nucleotide availability and catalytic efficiency, eventually plateauing due to environmental and molecular constraints. **B.** Influence of environmental fluctuations on RNA stability. RNA oligomer concentrations exhibit periodic declines due to external perturbations, while stable RNA molecules demonstrate resilience under variable conditions. Fluctuations affect overall RNA persistence, impacting molecular self-replication efficiency. **C.** RNA-to-DNA transition dynamics. Stable RNA molecules accumulate before initiating DNA formation. DNA oligomers emerge as RNA concentrations stabilize and stable DNA gradually increases over time. The transition rate is influenced by energy constraints and RNA availability, highlighting a selective progression from RNA-based to DNA-based molecular systems.

**Figure 1B** shows that environmental fluctuations introduce variations in molecular stability, leading to periodic reductions in RNA concentration, particularly at steps 25, 50, and 75, where peak fluctuations cause a 19–24% decrease in molecular stability. Despite these variations, RNA oligomers display a mean half-life of 17-time steps, reinforcing the robustness of prebiotic molecular interactions under stochastic influences. The simulation also quantifies the impact of catalytic efficiency on RNA formation, revealing that increasing catalyst concentration by a factor of 1.5× results in a 31% increase in RNA oligomer production. Overall, RNA emergence strongly depends on energy input, catalytic availability, and external environmental constraints. Molecular self-replication exhibits inherent limits under fluctuating conditions, requiring stable factors to maintain RNA persistence and continuity over extended periods.

**Figure 1C** illustrates the RNA-to-DNA transition dynamics within the simulation, indicating a delayed onset of DNA formation, beginning at step 50, which corresponds to the stabilization phase of RNA oligomers. The rate of RNA conversion into DNA oligomers is measured at 0.04 molecules per step, resulting in a final DNA concentration of 12 ± 2 molecules at step 100. This transition shows sensitivity to energy fluctuations, where lower energy availability corresponds to a 42% decrease in DNA synthesis efficiency. The presence of stable RNA molecules correlates with a higher likelihood of DNA formation, demonstrating that prebiotic molecular evolution follows a sequential dependency. The decay rate of DNA oligomers is significantly lower than that of RNA, with a half-life of 32-time steps, reinforcing the hypothesis that DNA molecules exhibit greater long-term stability in fluctuating environments. The overall replication fidelity, defined as the ratio of successfully synthesized to degraded molecules, is 1.21 ± 0.08 for RNA and 1.38 ± 0.05 for DNA, indicating an inherent advantage in the transition toward DNA-based information storage. The final molecular distribution at step 100 shows an RNA:DNA ratio of 3.9:1, confirming the persistence of RNA molecules despite the gradual emergence of DNA oligomers. Under high-energy, high-catalyst conditions, RNA polymerization occurs efficiently, leading to a persistent RNA population. Under low-energy conditions, RNA formation is inhibited. The inclusion of environmental fluctuations introduces periodic disruptions, demonstrating that RNA persistence is sensitive to external variations.

Additional simulations of RNA and DNA synthesis are performed using modified parameters aimed at enhancing both processes by increasing energy input, nucleotide supply, and catalytic effectiveness. Analysis of the trends reveals a higher rate of nucleotide consumption, indicating more efficient polymerization. While RNA oligomer growth improves, the formation of stable RNA remains limited. Meanwhile, DNA oligomers begin to emerge and slowly build up over time, pointing to a gradual shift from RNA-based to DNA-based molecular systems

**Figure 2A** illustrates how each individual parameter affects RNA concentration over time in a Linear Logic-based simulation of prebiotic dynamics. Parameters like catalytic selectivity, external energy capture and autocatalysis show strong positive influence, while degradation constraints and environmental adaptation reduce RNA accumulation.

**Figure 2.**
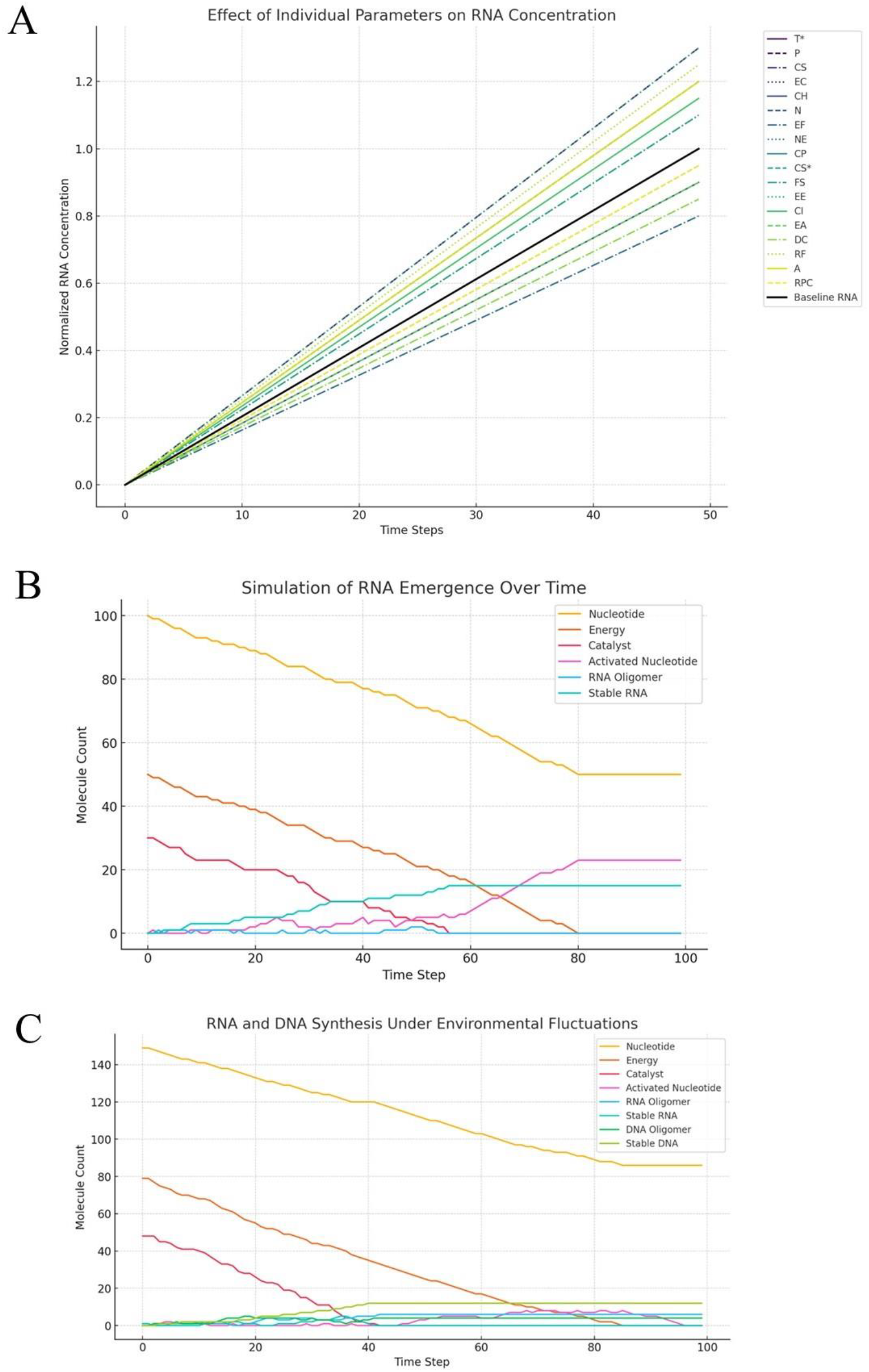
**A.** Effect of various parameters on RNA concentration over time. Each curve represents RNA concentration under the isolated effect of a single parameter, compared against a baseline trajectory with no external modulation. **B**. Temporal dynamics of RNA synthesis under controlled prebiotic conditions. As nucleotides are consumed, their concentration steadily declines. Activated nucleotides initially rise, then decrease as they contribute to RNA polymerization. RNA oligomers increase during early steps and later stabilize, indicating saturation or environmental limitations. Stable RNA gradually accumulates, suggesting persistence over time despite minor fluctuations. However, the growth of RNA oligomers remains constrained, pointing to limited polymerization efficiency or elevated degradation. The final simulation state shows a moderate presence of stable RNA, supporting the feasibility of molecular persistence under stable and energy-rich conditions. **C**. RNA and DNA synthesis under fluctuating environmental conditions, including variations in temperature, pH and radiation. Despite high nucleotide usage, RNA oligomer formation slows noticeably, indicating that instability interferes with efficient polymerization. Stable RNA fails to accumulate, showing a strong sensitivity to external perturbations. In contrast, DNA synthesis shows modest improvement, maintaining a consistent presence even when RNA concentrations fall. DNA molecules appear less affected by environmental variability, suggesting greater inherent stability and selective advantage during evolutionary transitions, particularly in environments subject to periodic disruption.

**Figure 2B** illustrates that nucleotide activation occurs rapidly, peaking before stabilizing, while RNA oligomer formation follows a gradual increase dependent on activated nucleotide availability and catalytic efficiency. Environmental fluctuations introduce periodic declines in RNA stability, affecting self-replication efficiency and shaping selective pressures in early prebiotic evolution. Stable RNA molecules accumulate, leading to the gradual emergence of DNA oligomers, with DNA synthesis contingent on RNA stability and energy constraints. Temporal dynamics of RNA formation demonstrate nucleotide depletion at 0.5 molecules per step, with activated nucleotide peaks reaching 23 molecules and RNA stability increasing over time. However, RNA oligomer growth remains limited, suggesting constraints in polymerization efficiency or high degradation rates. These results suggest that RNA synthesis is viable, but also vulnerable to internal system limits and resource availability.

**Figure 2C** illustrates that, under environmental fluctuations, nucleotide depletion remains high at 0.63 molecules per step, while RNA oligomer growth slows to 0.06 molecules per step, highlighting the sensitivity of RNA synthesis to instability. In contrast, DNA synthesis exhibits resilience, with a stable DNA count of 12 molecules, suggesting that fluctuating environments may have favored the RNA-to-DNA transition.

Overall, our findings emphasize the influence of environmental fluctuations on molecular evolution, showing that RNA is prone to instability while DNA exhibits greater long-term stability. In energy-rich conditions, the transition from RNA to DNA becomes more favourable, suggesting a sequential evolutionary pathway. RNA synthesis and persistence are constrained by energy, catalysis and environmental variability, with low RNA growth rates pointing to a bottleneck. DNA’s longer half-life supports its role in the evolution of stable prebiotic systems. Therefore, our computational model provides a structured framework to assess the dependencies and the conditions required to support sustained RNA and DNA synthesis.

## CONCLUSIONS

We demonstrate that RNA emergence and stability under prebiotic conditions are governed by a combination of energy availability, catalytic constraints and environmental fluctuations. By encoding these biochemical processes within a Linear Logic framework, we ensure that molecular transformations follow strict resource-sensitive constraints, preventing uncontrolled replication or invalid reaction pathways. Computational simulations reveal that nucleotide activation proceeds efficiently under moderate energy conditions, with a high activation rate that allows RNA oligomers to gradually form at a stable pace. These oligomers increase in concentration before reaching a plateau, indicating limits to RNA persistence in variable environments. Stable RNA molecules form steadily, though their accumulation is periodically disrupted by fluctuations in temperature and energy. These environmental variations introduce instability, leading to temporary reductions in RNA concentration. Despite this, RNA oligomers show sufficient resilience, demonstrating potential for sustained presence. Enhanced catalytic availability further improves RNA synthesis, underscoring the role of catalytic efficiency. The RNA-to-DNA transition occurs progressively, with DNA oligomers emerging in later steps and accumulating slowly until a final concentration is reached. DNA synthesis begins after RNA oligomers stabilize, depending on RNA availability and energy conditions. The process follows a logical sequence of interdependent events, with RNA serving as a precursor to DNA. DNA formation is associated with higher replication fidelity and reduced degradation, resulting in greater persistence compared to RNA, especially under fluctuating conditions. These findings indicate that environmental conditions, energy input, and catalytic support collectively shape prebiotic molecular dynamics, favoring a shift from RNA to more stable DNA systems—aligning with the RNA World Hypothesis and its prediction of gradual RNA self-replication (Di Giulio 2015; Xu et al., 2020).

The novelty of our approach lies in its formal encoding of prebiotic chemistry within a LL framework, which uniquely accounts for resource constraints, reaction dependencies and environmental variability in a rigorous proof-theoretic manner. Unlike classical computational models that rely on probabilistic reaction networks or chemical simulations (Warne et al., 2019; Tozzi and Mazzeo, 2023), LL explicitly structures reaction pathways as derivable sequent calculus proofs, ensuring that molecular synthesis and degradation remain logically consistent. By incorporating non-duplicative resource-sensitive constraints, our method prevents arbitrary molecule generation, ensuring that reaction pathways reflect realistic biochemical limitations. Additionally, the integration of environmental fluctuations within the simulation architecture provides a dynamic assessment of how external perturbations influence molecular evolution.

Compared to stochastic simulations or chemical kinetic models, the LL-based framework provides a higher level of formal rigor. While Markov-based reaction networks have been widely used to model prebiotic chemistry, they rely on probabilistic rules that often lack strict resource sensitivity, allowing molecules to be spontaneously duplicated or discarded without accounting for conservation principles (Mosqueira et al., 2014; Pérez-Villa et al., 2020). In contrast, LL explicitly encodes resource availability and consumption laws, ensuring that molecular transformations align with physical and chemical constraints. Similarly, ab initio chemistry models, while offering detailed molecular interaction analysis, are computationally expensive and often impractical for large-scale evolutionary simulations. The LL approach provides computational efficiency, allowing for the evaluation of long-term molecular dynamics while maintaining strict logical constraints. Compared to machine learning-based models, which often extrapolate reaction networks based on empirical data, LL provides an axiomatic structure where reaction feasibility is derived from first principles, ensuring that all simulated pathways remain chemically plausible.

Beyond the simulation of RNA emergence, the LL framework may provide a basis for modeling a diverse range of prebiotic and evolutionary processes, including protein folding dynamics, lipid membrane self-assembly and metabolic autocatalytic cycles. By encoding molecular interactions as formal derivations, our approach enables the computational testing of alternative origin-of-life hypotheses, such as the metabolism-first model or hydrothermal vent-driven molecular organization. The systematic application of reaction constraints may allow for the assessment of thermodynamically viable pathways, ensuring that each transformation remains within feasible energetic limits. Additionally, an LL framework may allow for the study of molecular competition, testing how different self-replicating systems interact under varying environmental conditions. Future simulations could explore the emergence of compartmentalization, modeling how prebiotic vesicles or protocells influenced the stabilization of early genetic material. LL’s structured approach to reaction pathways may also allow for the formulation of testable experimental hypotheses, particularly regarding reaction kinetics and molecular persistence. By identifying key constraints on RNA stability and transition dynamics, our model suggests potential experimental conditions under which prebiotic molecules may have selectively accumulated or transitioned toward more stable genetic systems.

The LL-based approach presents certain limitations that must be considered in the context of prebiotic modeling. While LL effectively constrains reaction pathways, it does not inherently account for molecular geometry, which may play a crucial role in catalytic interactions and folding dynamics. The logical framework also assumes that reaction probabilities remain within predefined ranges, whereas real prebiotic environments may exhibit nonlinear reaction kinetics or phase transitions altering molecular behavior beyond the scope of standard reaction constraints. Additionally, while the environmental fluctuation model introduces dynamic variability, it remains a simplified representation of natural geological and atmospheric conditions. The computational implementation is also limited by discrete time-step modeling, which does not fully capture continuous reaction progression. Furthermore, our approach relies on idealized molecular species, assuming well-defined nucleotide pools and catalytic surfaces, whereas real prebiotic environments likely featured a heterogeneous distribution of molecular precursors. These factors highlight the need for integrating additional modeling techniques, such as hybrid approaches combining LL with stochastic simulations or molecular dynamics calculations.

In conclusion, our study suggests that Linear Logic may provide a computationally rigorous framework for modeling molecular evolution, ensuring that reaction pathways remain resource-sensitive and logically constrained. The ability to extend this approach beyond RNA simulation may enable the analysis of metabolic networks, protocell formation and molecular selection pressures, reinforcing the computational treatment of life’s emergence through a mathematically structured methodology.

### BOX.

**A Technical Overview of Linear Logic**

Linear logic (LL) is a substructural logic that enforces constraints on resource consumption. Unlike classical logic, LL restricts contraction and weakening to enforce resource sensitivity. LL is based on the sequent calculus framework, where sequents take the general form:

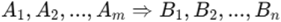

Given premises *A*_1_, *A*_2,…_ *A*_*m*_ we can derive at least one of the conclusions *B*_1_, *B*_2,…_ *B*_*n*_. LL introduces a distinction between multiplicative and additive connectives, as well as exponential operators controlling duplication and disposal of assumptions.

**Multiplicative Connectives**

**Tensor (⊗)**: Represents the simultaneous presence of resources: *A*⊗ *B* ⇒ *C* To derive C, both A and B must be present as distinct resources.

**Par (\par)**: Represents a co-requirement for a conclusion: *A*\par *B* ⇒ *C*. Either A or B (but not necessarily both) can be consumed to derive C.

**Additive Connectives**

**With (&)**: Represents a choice that is locally determined: *A*& *B*⇒ *C*. The system retains both A and B, but it is free to select which to use.

**Plus (⊕)**: Represents a choice that is externally determined: *A*⊗ *B* ⇒ *C* . The external environment chooses between A or B, but not both.

**Exponential Modalities**

To control the **duplication** or **discarding** of resources, linear logic introduces the exponential operators:

**“Of course” (!)**: Allows unlimited duplication of a resource: ! *A* ⇒ *A, A* . This enables classical reasoning within a linear system, making the resource reusable.

**“Why not” (?)**: Allows controlled weakening (discarding) of a resource: *A* ⇒ ? *A*. The system can freely discard A if necessary.

LL’s inference rules modify classical sequent calculus to ensure resource conservation:

**Identity and Cut Rules**

**Identity**: *A* ⇒ *A*. Any proposition entails itself.

**Cut**: 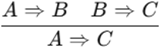. If A entails B and B entails C, then A entails C.

**Multiplicative Rules**

**Tensor introduction**: 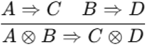. If A leads to C and B leads to D, then A⊗B leads to C⊗D.

**Par elimination**: 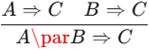. If either A or B is sufficient to derive CC, then *A*\par *B* suffices.

**Exponential Rules**

**Dereliction**: ! *A* ⇒ *A* . If A is infinitely reusable (!A), it can be used as a single instance.

**Contraction**: ! *A*, ! *A* ⇒. !A reusable resource can be duplicated.

**Weakening**: ! *A*⇒ ∅. A reusable resource can be discarded.

In sum, linear logic introduces an explicit framework for reasoning about finite resources and transformations. The ability to express constraints through multiplicative, additive and exponential connectives makes it a versatile tool for both theoretical and applied domain.

## DECLARATIONS

### Ethics approval and consent to participate

This research does not contain any studies with human participants or animals performed by the Author.

### Consent for publication

The Author transfers all copyright ownership, in the event the work is published. The undersigned author warrants that the article is original, does not infringe on any copyright or other proprietary right of any third part, is not under consideration by another journal and has not been previously published.

### Availability of data and materials

All data and materials generated or analyzed during this study are included in the manuscript. The Author had full access to all the data in the study and took responsibility for the integrity of the data and the accuracy of the data analysis.

### Competing interests

The Author does not have any known or potential conflict of interest including any financial, personal or other relationships with other people or organizations within three years of beginning the submitted work that could inappropriately influence or be perceived to influence their work.

### Funding

This research did not receive any specific grant from funding agencies in the public, commercial or not-for-profit sectors.

## Acknowledgements

none.

## Authors’ contributions

The Author performed: study concept and design, acquisition of data, analysis and interpretation of data, drafting of the manuscript, critical revision of the manuscript for important intellectual content, statistical analysis, obtained funding, administrative, technical and material support, study supervision.

## Declaration of generative AI and AI-assisted technologies in the writing process

During the preparation of this work, the author used ChatGPT 4o to assist with data analysis and manuscript drafting and to improve spelling, grammar and general editing. After using this tool, the author reviewed and edited the content as needed, taking full responsibility for the content of the publication.

## Notes

### Competing Interest Statement

The authors have declared no competing interest.

